# Ventral Pallidum Cholinergic Neurons Respond to Reward and Signal Reward Value

**DOI:** 10.64898/2026.05.24.726981

**Authors:** Alexander Gannon, Ruby Setara, Czarina M. Maysonet, Ronald E. Salazar, Dustin R. Zuelke, Jane E. Bowen, Eduardo F. Gallo

**Affiliations:** Department of Biological Sciences, Fordham University, Bronx, NY

## Abstract

The ventral pallidum (VP) is a key basal ganglia structure involved in integrating reward-related signals to guide motivated behavior and feeding. VP neuronal activity is strongly tuned to reward-predictive cues and palatable rewards, tracking changes in value and motivational state. Recent studies have established important and opposing roles for GABAergic and glutamatergic VP neurons in reward- and avoidance-related behaviors. However, much less is known about the role of VP cholinergic neurons (VP-CNs), which comprise only ∼10% of VP neurons in mice, but maintain extensive connectivity with reward-related circuitry. Here, using cell type-selective *in vivo* fiber photometry during instrumental reward behavior, we found that VP-CN activity responds to cues, actions, and reward retrieval in a training-independent manner. Phasic VP-CN responses to reward retrieval were reduced by reward dilution, whereas pre-feeding with the reward enhanced these responses. In contrast, non-cholinergic VP neurons exhibited modulation by pre-feeding but not by reward dilution. Together, these findings identify VP-CNs as a distinct and dynamically regulated VP population whose activity is sensitive to both reward palatability and physiological state, extending our current understanding of the VP and its roles in reward processing.

## INTRODUCTION

Reward processing enables individuals to respond to rewards and update value based on changing reward properties, costs, environmental cues, and physiological state [1]. These functions, which are critical for motivation and decision-making, engage a broad network of brain areas, including the ventral pallidum (VP), a key node of the ventral basal ganglia [2,3]. The VP’s connectivity to areas critical to motivated and consummatory behaviors like nucleus accumbens (NAc), basolateral amygdala (BLA), prefrontal cortex (PFC), ventral tegmental area (VTA), and hypothalamus has prompted numerous efforts to define the VP’s role in integrating distinct reward-related signals and mediating adaptive behavioral output [3–8].

Extensive work has highlighted the VP’s contributions to different aspects of motivation for palatable food rewards. For instance, VP neuron firing encodes vigor of reward-seeking actions, while VP inactivation reduces effort expenditure for food [9–12]. The VP also plays important roles in the processing of reward information. *In vivo* recordings demonstrated that most VP neurons were consistently activated by sucrose reward consumption, regardless of cue-reward learning [13]. “Hedonic hotspots” of VP neural activity track the enhanced orofacial reactions elicited by palatable rewards [14]. Moreover, VP activity tracks the expected or relative value of rewards and is sensitive to manipulations that alter reward size, appetite, or that pair reward with unpleasant stimuli [14–18]. Ablation and pharmacological inactivation of the VP not only suppress food consumption but also produce aversive reactions and decrease motivation toward palatable rewards [19–23]. Ample evidence across species has also revealed that VP activity is highly sensitive to presentation of reward-related cues [11,13,24–26] and mediates cue-induced invigoration and reinstatement of reward seeking [11,27,28].

Circuit-level studies have expanded these findings by demonstrating that GABAergic VP neurons, which account for nearly 75% of VP neurons, are critical to positive valence reward behavior [23,29–33]. In contrast, VP glutamatergic neurons, a minority sub-population, have been recently linked to negative value encoding and avoidance [30,34]. However, less is known about the reward-related contributions of cholinergic neurons in the VP (VP-CNs), which constitute approximately 10% of VP neurons in mice and distinctly innervate key reward structures like the BLA and PFC [29,35]. Recent work reported that VP-CNs are necessary for approach to appetitive olfactory stimuli [36] and that activation of VP-CN terminals in the BLA increases breakpoint in a progressive ratio task [37]. In addition, VP-CNs respond to reinforcement-related cues and outcomes in a Pavlovian task [38]. However, it remains unclear whether VP-CN activity is modulated by changes in reward value or physiological state during instrumental reward seeking.

Here, using *in vivo* fiber photometry to monitor calcium dynamics in VP-CNs during a continuous reinforcement (CRF) task in freely moving mice, we demonstrate that VP-CN activity robustly increases in response to a palatable food reward and is dynamically modulated by changes in reward value and satiety state. These results identify VP-CNs as a distinct component of ventral pallidal reward processing.

## MATERIALS AND METHODS

### Animals

Adult male and female homozygous ChAT-IRES-Cre mice (ChAT*^tm1(cre)Lowl^/MwarJ;* JAX stock #031661)[39], were bred in-house and used for all experiments. Mice (8-10 months of age) were housed in groups of 3-5, when possible, and kept on a 12-h light/dark cycle, and all experiments were performed during the light cycle. All experimental procedures were conducted following NIH guidelines and were approved by Institutional Animal Care and Use Committee of Fordham University.

### Surgery

Mice underwent stereotaxic surgical procedures under isoflurane anesthesia to receive unilateral infusions (180 nL) of adeno-associated viruses (AAVs) into the VP using the following coordinates at AP, +0.26; ML, +1.5; and three DV sites: -4.45, -4.55, -4.65 from skull surface) at a rate of 20 nl/s (9 pulses, 5 min). Separate cohorts of mice received either AAV9-syn-FLEX-jGCaMP7f-WPRE (Addgene viral prep # 104492-AAV9, 2.2 x 10^13^ gc/mL)[40] or a Cre-Off AAV1-hSyn-FAS-GCaMP6f [41] (a gift from Christoph Kellendonk, produced at Vector Biolabs, 8.0 x 10^13^ gc/mL). Following virus injection, a fiber optic cannula (400-μm core, 0.66 N.A., Doric, Quebec, Canada) was inserted through the same opening (AP: +0.26, ML: +1.5, DV: -4.55 from skull surface) and secured to the skull using dental cement anchored to stainless steel microscrews.

### Behavior

Operant chambers (Env-307w; Med-Associates, St. Albans, VT) equipped with liquid dippers inside light- and sound-attenuating cabinets were used. Chamber interior flooring consisted of metal rods spaced 0.87 cm apart and an open ceiling. One wall of the chamber contained a central feeder trough. Raising of the dipper arm delivered a drop (13-15 μL) of undiluted evaporated milk. Head entries into the trough were recorded with an infrared (IR) photocell detector. Two retractable levers were mounted on either side of the trough. A house light located opposite the trough illuminated the chamber during behavioral sessions. The experimental protocols were controlled via Med-PC computer interface and Med-PC V software.

Mice initiated behavioral training 3 weeks after surgery. Operant tasks were conducted under mild food restriction (85-90% of baseline body weight) with water available *ad libitum*. During the initial dipper training session, 20 dipper presentations were delivered on a variable inter-trial interval (ITI), and sessions ended after 20 rewards were retrieved or after 30 min had elapsed, whichever occurred first. Criterion consisted of head entries during all 20 dipper presentations within a single session. In the second dipper training phase, mice were fit with patch cords, and criterion was achieved when all 20 rewards were retrieved within 30 min.

For lever press training, presses on a single lever were immediately reinforced with a milk reward on a continuous reinforcement (CRF) schedule. Mice were randomly assigned to one of two levers and remained on the same lever throughout the experiment. CRF trials began with lever extension, and levers were retracted 5 s after the initial press before being presented again after a variable ITI (average 20 s). The reward consisted of a 5-s dipper presentation.

Initially, mice were trained without patch cords until 25 dippers were earned in a 30-min CRF session. Mice were then fitted with patch cords for recording during CRF-1s schedule, in which rewards were delivered 1 s after the first lever press. These sessions lasted for either 60 minutes or until 60 rewards were earned, whichever occurred first. Fiber photometry data were collected from the first four CRF-1s sessions following the initial no-patch cord session. To allow greater temporal separation of behavior events during dilution and pre-feeding studies, mice were trained for 3 days on CRF-2s, in which the reward delivery was delayed by 2 s following the first press.

Mice then underwent alternating CRF-2s sessions of pre-feeding and no pre-feeding for 4 days. For pre-feeding, mice were placed into individual cages (26 cm x 15 cm) equipped with a 50-ml polycarbonate tube fitted with a ball point stainless steel sipper tube for free access to evaporated milk for 30 minutes prior to the session. Milk consumed was recorded by weighing tubes before and after pre-feeding. For dilution experiments, the same mice underwent six CRF-2 sessions in which milk rewards were diluted with water and presented in the following concentration order across days: 100%, 10%, 25%, 2%, 50%, and 0%. All mice received the same reward concentration on a given day.

Behavioral data recorded from MedPC was analyzed using custom Python scripts. Rewards earned were measured as the number of dippers presented following lever presses, while retrievals were the number of dipper presentations which coincided with head entries in the feeder trough, as measured by IR beam breaks. Trials were labeled as retrieved if they included a detected head entry during the five seconds of reward presentation. Press latency was calculated as the time between lever extension and the first press. Trials with “long” press-latency were those with more than 3 s between lever extensions (lever on) and lever press.

Head poke latency was the time between dipper presentation and entry into the feeder trough, while head poke duration was the total head poke time during the 5-s dipper presentation.

### In Vivo Fiber Photometry and Analysis

Fiber photometry (FP) recordings were performed throughout behavior sessions using a TDT LUX RZ-10X processor and Synapse software (Tucker-Davis Technologies) with real-time lock-in amplification to isolate fluorescence signals from background noise. Excitation light at 465 nm and 405 nm was delivered via fiber-coupled LEDs and sinusoidally modulated at distinct frequencies (210 and 330 Hz). Light power at the fiber tip was maintained at approximately 55 µW (465 nm) and 10 µW (405 nm). Light was routed through a fluorescence iFMC6 mini-cube and low-autofluorescence patch cords (Doric Lenses), which were photobleached prior to use to minimize baseline fluorescence and coupled to an implanted optical fiber cannula. Emitted fluorescence was collected through the same optical path, demodulated, low-pass filtered at 6 Hz, and recorded at 1017 Hz.

FP data were preprocessed using GuPPy [42]. To account for motion artifacts and photobleaching, the 405 nm control channel was fit to the 465 nm signal channel using least-squares linear regression. The fitted 405 nm signal was then subtracted from the 465 nm signal to generate ΔF/F values. Resulting signals were smoothed using a moving average filter (0.1-s window) and z-scored across the recording session. Behavioral events were generated by Med Associates software and recorded as TTL pulses in the TDT system, providing shared timestamps with photometry data. Peri-event time histograms (PETHs) were generated by aligning z-scored signals to behavioral event onset and extracting activity within a defined time window surrounding each event. Signals were averaged across trials within each subject prior to group-level analyses for each session. Behavioral events included lever extension, lever press, dipper delivery, and rewarded head entry (HE). Rewarded HE was defined as the first instance during the 5-s reward presentation in which the mouse was detected in the feeder trough.

Peak amplitudes and area under the curve (AUC) were calculated from session-averaged data using custom Python scripts. Peak amplitude was defined as the maximum value within a 0-1 s post-event window and was used to quantify phasic responses aligned to Lever On and Lever Press events. AUC was computed over the same interval using the trapezoidal method. For Rewarded HE, both peak amplitude and AUC within 0-1s were quantified to capture both the maximal and overall magnitude of reward retrieval–related activity during the immediate post-entry epoch, while minimizing contributions from variable head entry duration and subsequent behaviors. For each animal, a minimum of 5 trials per event was required for inclusion in averaged traces and for calculation of peak amplitude and AUC.

### Immunofluorescence

Mice were transcardially perfused with ice-cold 4% paraformaldehyde (Fisher Scientific, Pittsburgh, PA) in PBS under deep anesthesia (100 mg/kg ketamine and 10 mg/kg xylazine, i.p.). Fixed brains were harvested, postfixed overnight, and washed in PBS before 50-μm coronal sections were prepared with a Leica VT1200S vibratome (Richmond, VA). After a 2-h incubation at room temperature in blocking solution (10% fetal bovine serum, 0.5% Triton X, 0.5% bovine serum albumin in TBS), sections were labeled overnight at 4°C with primary antibodies against ChAT (goat; 1:100, AB144P Millipore Sigma, Burlington, MA) and GFP (chicken; 1:1000; AB13970 Abcam, Cambridge, MA). Sections were incubated with corresponding fluorescent secondary antibodies for 2 h at room temperature, washed, mounted on slides, which were coverslipped with Vectashield Hardset with DAPI (Vector, Burlingame, CA). Images were obtained with a Nikon Eclipse Ti-E epifluorescence microscope or a Leica TCS SP8 laser-scanning confocal microscope and processed with NIH Image J software. The position of implanted fibers was estimated by manual alignment to the Franklin and Paxinos brain atlas (3^rd^ edition).

### Data Analysis

Statistical analyses were performed using GraphPad Prism 10 (GraphPad). Data are expressed as mean ± the standard error of the mean (SEM). Paired two-tailed Student’s t-tests were used to compare 2-group data. Multiple comparisons were analyzed using one- or two-way repeated-measures ANOVA or mixed-effects models, as appropriate. A p-value of < 0.05 was considered statistically significant.

## RESULTS

### VP-CN activity increases in response to cues, lever presses and rewards

To determine whether VP-CNs respond during reward-related behavior, we used *in vivo* fiber photometry (FP) analysis to monitor bulk Ca^2+^ activity from VP-CNs. To target VP-CNs, we injected a Cre-dependent adenoassociated virus (AAV) expressing GCaMP and implanted an optic fiber into the VP of ChAT-IRES-Cre mice. This achieved selective co-expression of GCaMP with choline acetyltransferase (ChAT), a marker of cholinergic neurons **(Fig. 1A-B)**. In addition, GCaMP positive terminal fields were found in the BLA of ChAT-IRES-Cre mice **(Fig. 1C, D)**, but not Cre-negative mice **(Fig. 1E, F)**, consistent with the known projections from VP-CNs to BLA [5,35]. Fiber implant placement within the VP was histologically verified following behavioral experiments **(Fig. 1G, H)**.

**Figure 1.**
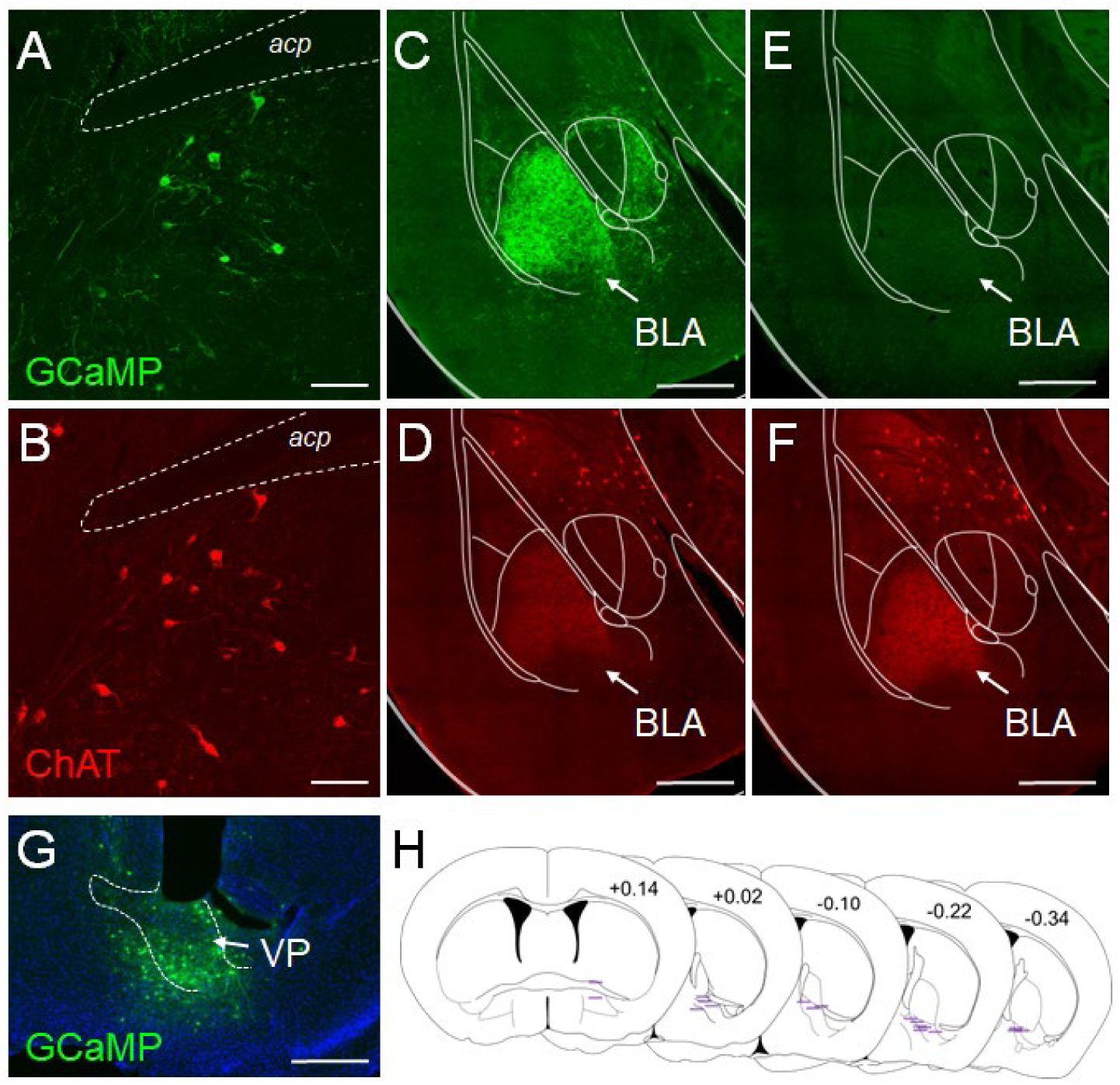
Cell-selective targeting of VP-CNs in ChAT-IRES-Cre mice. **A, B.** Co-immunolabeling of AAV9-Flex-jGCaMP7 (GFP, *green*) and the cholinergic marker ChAT (*red*) shows specific GCaMP expression in VP-CNs of ChAT-IRES-Cre mouse 4 weeks after injection. Scale bar, 100 μm. *acp* = anterior commissure. GCaMP-positive VP-CN terminals found in the BLA of a ChAT-IRES-Cre mouse (**C, D**) but not in a Cre-negative control (**E, F**) receiving AAV9-Flex-GCaMP7 in VP. Scale bar, 400 μm. **G**. Representative image of GCaMP expression and fiber location in VP. Scale bar, 400 μm **H.** Summary of fiber optic cannula placements for experimental mice (n = 18), projected onto schematic coronal sections from the Paxinos and Franklin mouse brain atlas. Purple lines represent the estimated location of individual fiber tips, identified as the ventral-most point of the fiber track across serial histological sections. Numbers indicate distance (mm) from Bregma.

*In vivo* Ca²⁺ signals were recorded during continuous reinforcement (CRF) sessions consisting of 60 lever presentations delivered on a variable inter-trial interval (ITI) schedule. Each press resulted in delivery of a milk reward (one second after press, CRF-1s), followed by lever retraction. This short delay enabled analysis of VP-CN Ca^2+^ activity in response to temporally discrete task events: lever extension, lever press and reward presentation. To identify training-related effects on VP-CN responses to these events, we first assessed VP-CN activity across four consecutive sessions of early CRF training. We found a significant effect of training on CRF behavioral performance as reflected by decreased press latency following lever extension, increased rewards earned and rewards retrieved, decreased latency to retrieve rewards, and increased head entry duration during a reward presentation **(Fig. 2A-E)**.

**Figure 2.**
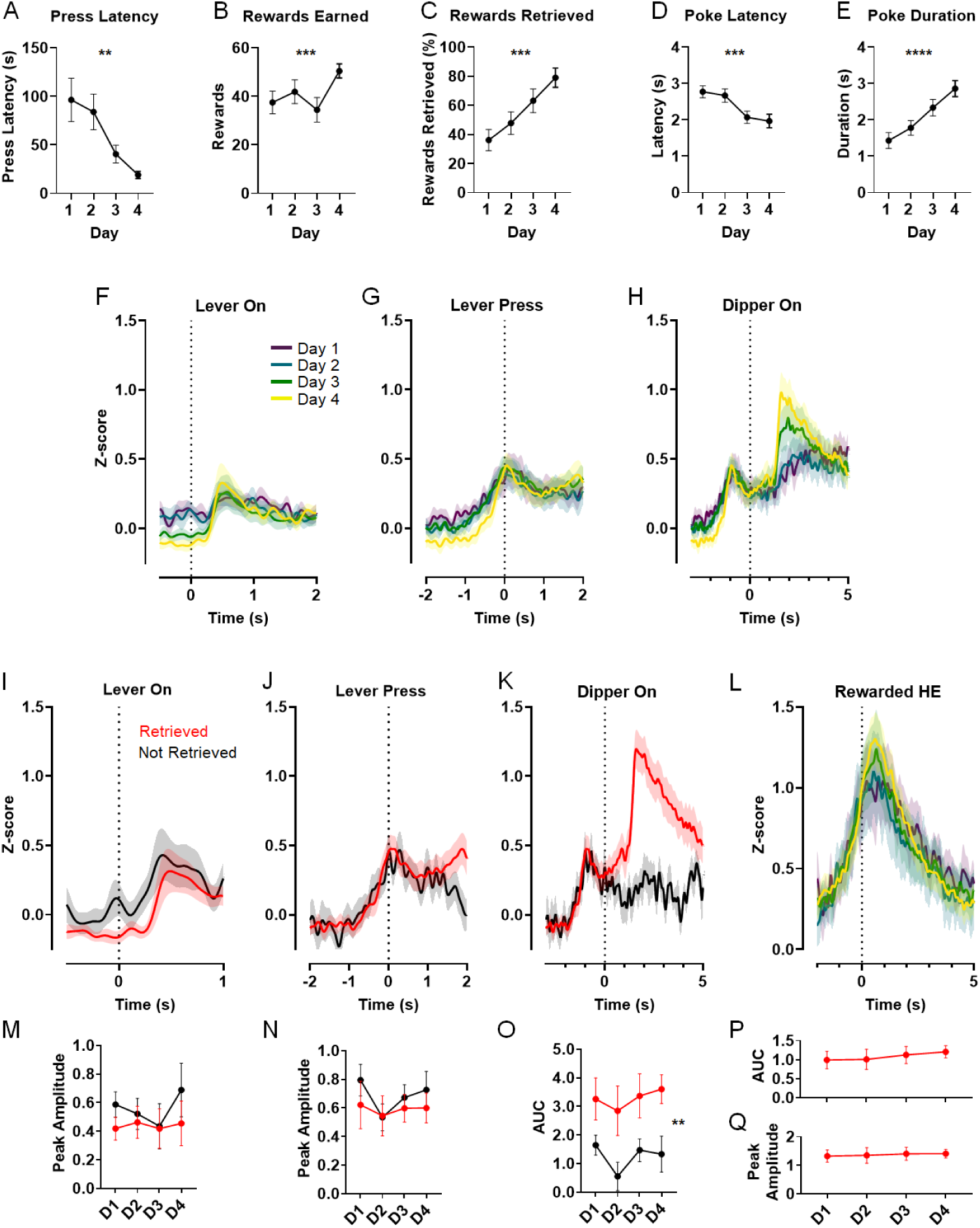
VP-CN activity increases in response to cues, lever pressing and reward. **A-E**. Across four CRF sessions where reward was delivered 1 s after a single press (CRF-1s), mice showed significant improvements in task performance, including increased rewards earned and retrieved (***p < 0.0005), and greater time spent in the feeder port during reward presentation (****p < 0.0001). Press latency and head entry/poke latency were also significantly reduced (**p <0.01, ***p< 0.0005). **F-H**. Mean z-scored GCaMP fluorescence aligned to lever extension (Lever On), lever press, and dipper presentation (Dipper On) across the first days of training. Dotted line denotes behavior event onset. **I-K**. Comparison of event-aligned mean GCaMP activity measured on Day 4, sorted by whether the reward was retrieved in each trial. **L**. Mean GCaMP activity across training days aligned to the first rewarded head entry (Rewarded HE) in each trial. **M, N**. Peak amplitude was not altered by training day or retrieval when aligned to Lever ON (Day: F _(3, 76)_ = 0.7192, p = 0.5436; Retrieval: (F_(1, 32)_ = 1.534, p = 0.2245) or Lever Press (Day: F _(3, 77)_ = 0.7324, p = 0.5358; Retrieval: (F_(1, 32)_ = 0.7564, p = 0.3909). **O**. Dipper On increased VP-CN activity in retrieved compared to non-retrieved trials (main effect of retrieval: F_(1, 32)_ = 9.603, **p = 0.004), independent of training day (F _(3, 76)_ = 1.110, p = 0.3502). **P, Q**. Neither A.U.C. (p = 0.1704) or peak amplitude (p = 0.6115) in response to Rewarded HE were affected by training. Data shown as mean ± SEM. n = 18 mice.

Lever extension, which becomes a predictive cue with training [43,44], elicited a small but consistent upward deflection in the average Ca^2+^ signal relative to the session baseline **(Fig. 2F)**. Similarly, VP-CNs exhibited increased phasic activity to the first lever press in each trial **(Fig. 2G)**. Peak responses within the 0-1 s post-event window showed no clear effect of training for either event **(Fig. 2M, N)**. The most robust activation of VP-CN signals occurred following onset of the 5-s reward presentation (Dipper On), which was especially prominent later in training **(Fig. 2H)**. Because reward retrieval efficiency also increased across sessions **(Fig. 2C)**, we sought to determine whether the Dipper On-related VP-CN activation was related to reward retrieval or to reward presentation alone. To this end, we sorted FP trial data based on whether the animal entered the reward port during reward presentation. While peak responses to lever extension or lever press did not differ based on retrieval **(Fig. 2I-J, M-N)**, VP-CN activation following Dipper On (0-5 s) was only observed in retrieved trials and did not vary across the 4 training sessions **(Fig. 2K, O)**. To isolate neural activity associated with initial reward retrieval, FP responses were aligned to the first rewarded head entry (HE) in each trial, and peak amplitude and AUC were quantified within a 0-1 s post-entry window **(Fig. 2L, P-Q)**. We found a similar degree of VP-CN activation following reward retrieval in all 4 sessions. These responses were comparable between males and females **(Fig. S1)**. These results show that while VP-CN activity is recruited by cues and instrumental actions, it responds most robustly to retrieval of the primary unconditioned reward, independent of training.

### VP-CN activity is modulated by reward palatability

We next sought to determine whether this reward-related activity is responsive to changes in the intrinsic properties of the reward. To address this question, we examined behavioral performance and VP-CN activity across six CRF sessions in which the milk reward was diluted with water to reduce its palatability. We first increased the delay between the lever press and reward presentation from 1 to 2 s (CRF-2s) to allow for a clearer distinction between lever press and reward-related neural activity. We confirmed stable behavior and neural activity across CRF-2s sessions **(Fig. S2).** Diluted milk (100, 50, 25, 10, 2, and 0%) was then presented to all animals in a fixed, pseudorandomized order across sessions. As expected, diluting the reward significantly increased press and head poke latency and decreased head poke duration and rewards earned and retrieved **(Fig. 3A-E)**. We also found that VP-CN activity evoked by reward retrieval decreased with diluted milk concentrations **(Fig. 3F)**. One-way ANOVA showed a significant reduction in both the AUC and peak amplitude at the onset of the reward retrieval response **(Fig. 3G,H)**. Despite no overall effect of dilution on lever extension signals **(Fig. 3I, J)**, significant differences emerged for Lever On in trials with long press latencies, during which the lever extension was more likely to be examined separately from the lever press **(Fig. 3K)**.

**Figure 3.**
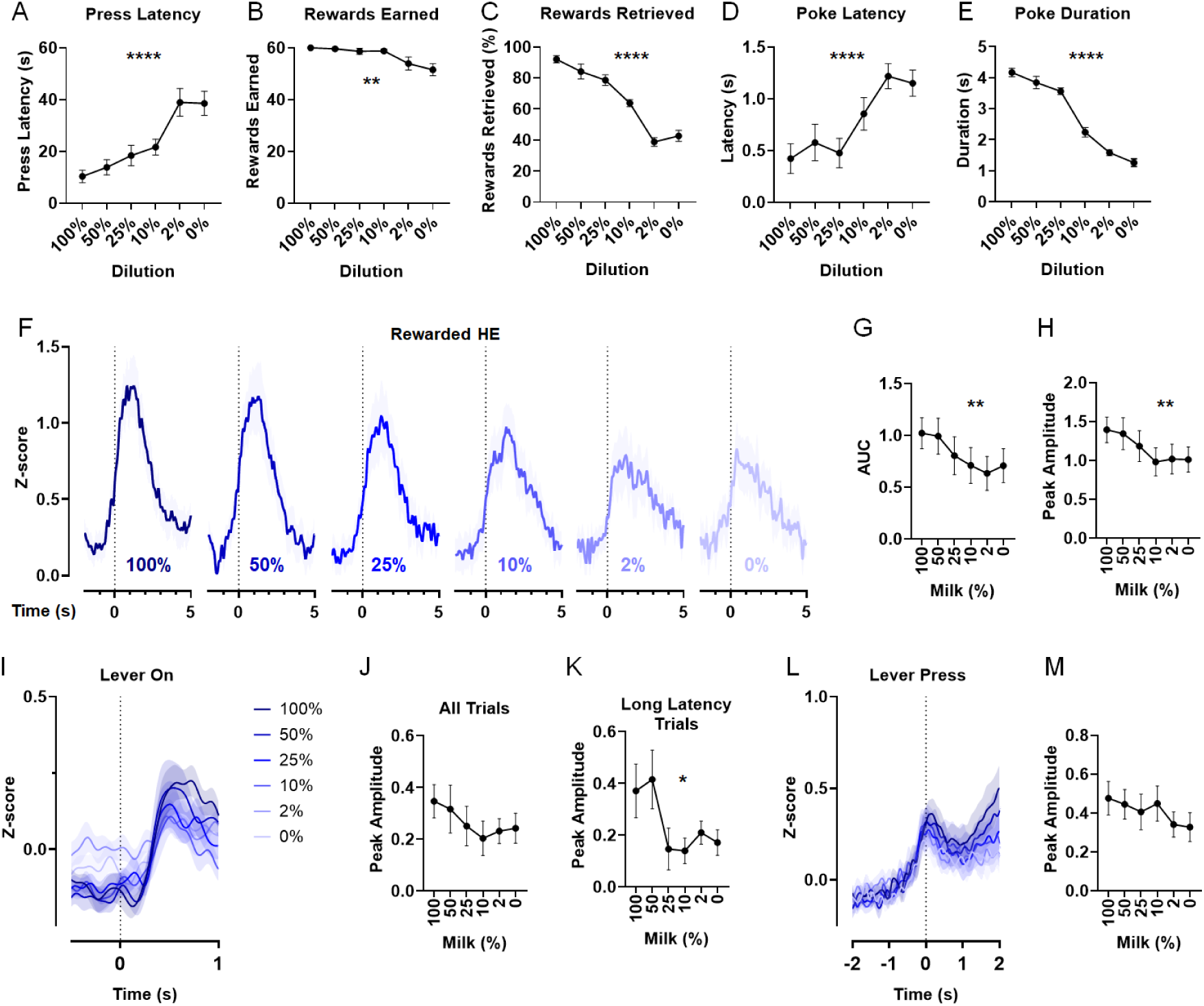
VP-CN Activity Tracks Reward Palatability. **A-E.** Behavioral performance across CRF-2s sessions as a function of milk reward concentration. RM one-way ANOVA indicated a significant effect of dilution on all measures. ** p<0.01, ****p <0.0001. **F.** Mean z-scored GCAMP signals time-locked to Rewarded HE at each milk concentration. **G, H.** A.U.C. and peak amplitude were significantly decreased as a function of dilution (A.U.C., F _(3.508, 42.10)_ = 4.827, **p = 0.0038, Peak, F _(3.576, 42.91)_ = 4.837, **p = 0.0035). **I, J.** Mean activity aligned to Lever ON across all trials did not show significant effects of dilution on peak amplitude F _(3.495, 41.93)_ = 1.172, p = 0.3347). **K.** However, in Lever On trials where the latency to press was longer than 3 s, dilution significantly reduced peak amplitude (F _(2.746, 32.40)_ = 3.336, *p = 0.0348). **L, M.** Lever press-related activity was not significantly altered by reward dilution (F _(3.236, 38.83)_ = 1.353, p = 0.2708). Data shown as mean ± SEM. n = 13 mice.

Furthermore, no significant dilution effects were observed for VP-CN responses to lever presses **(Fig. 3L,M)**. Together, these results suggest that VP-CN activity at reward retrieval and during the anticipatory phase is modulated by reward palatability.

### Pre-feeding-induced satiety increases the VP-CN reward response

Because reward value is influenced not only by the sensory properties of the reward but also by physiological state, we next assessed whether inducing satiety by pre-feeding modulates VP-CN reward responses. Mice were allowed to freely consume the milk reward for 30 min prior to two non-consecutive CRF-2s sessions. Within-subject comparisons showed a significant impact of pre-feeding (PF) across average behavioral measures relative to the no pre-feeding (NPF) sessions, indicating effective satiety-induced motivational drive reduction **(Fig. 4A-E)**. Notably, we found that phasic VP-CN Ca^2+^ activity in response to reward retrieval was significantly increased in PF compared to NPF sessions **(Fig. 4F-H)**. In contrast, PF did not alter peak responses to the lever press **(Fig. 4J)** or the lever extension **(Fig 4I)**, even in trials with long press latency (not shown). These findings indicate that pre-feeding-induced satiety specifically enhances phasic VP-CN responding to reward but not to cues or actions and suggests that reward-evoked VP-CN activity is modulated by physiological state.

**Figure 4.**
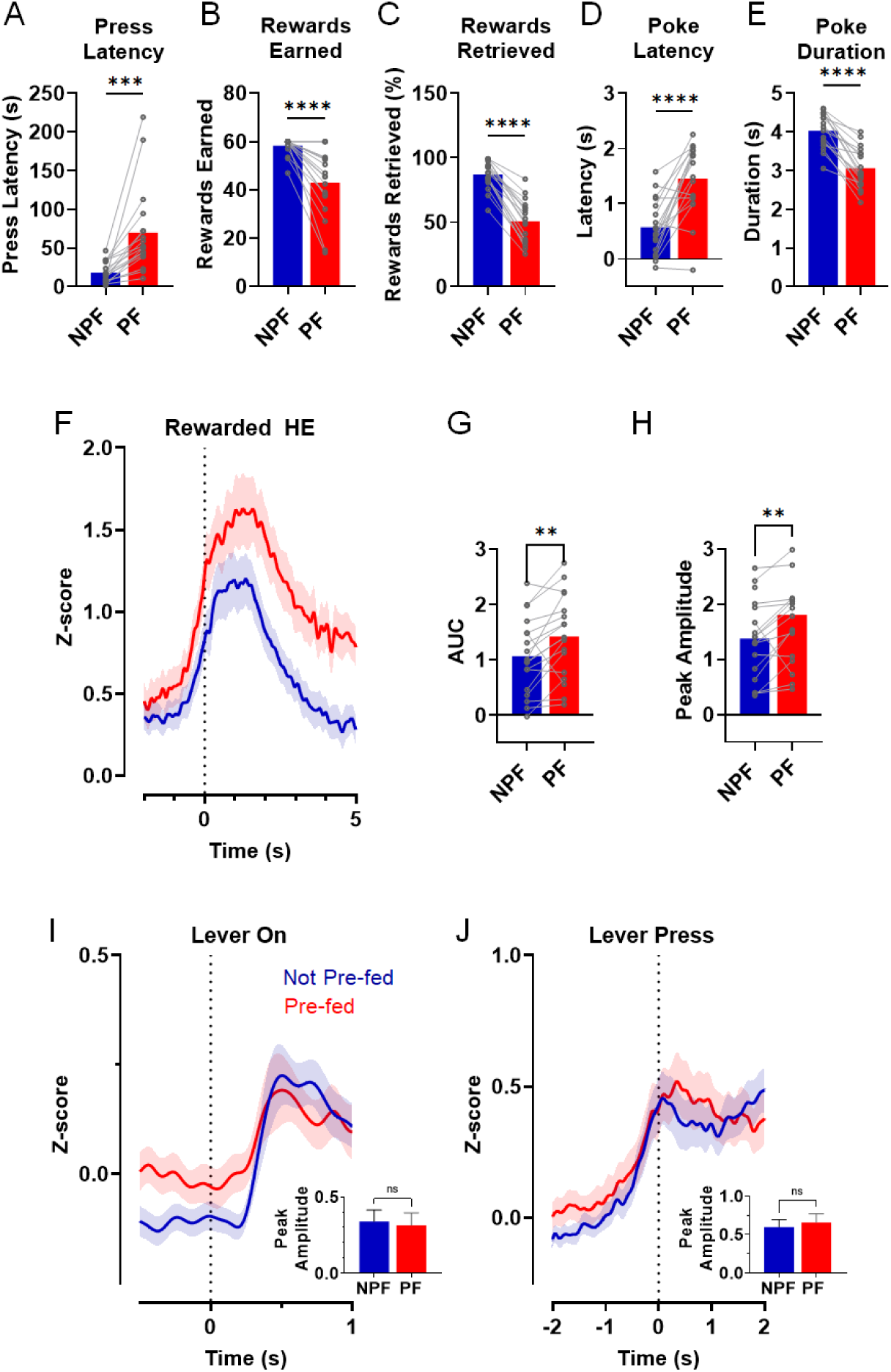
Pre-feeding-induced satiety increases the VP-CN reward response. **A-E.** Two-day average performance in CRF-2s sessions with and without pre-feeding with the milk reward (PF vs NPF). Paired t-tests revealed significant increases in press latency and poke latency, and significant reductions in rewards earned, rewards retrieved, and in poke duration (***p < 0.001; ****p<0.0001). **F.** Mean z-scored GCAMP responses time-locked to Rewarded HE in NPF vs PF sessions. **G, H**. Rewarded HE A.U.C. and peak amplitude were significantly increased following PF (**p < 0.01). **I, J.** GCaMP responses to Lever On (p = 0.65) or Lever Press (p = 0.2012) were not significantly altered by PF. Data shown as mean ± SEM. n = 18 mice.

### Non-cholinergic VP neuron reward responses are sensitive to satiety but not reward palatability

Because VP-CN reward responses were sensitive to both reward dilution and physiological state, we next examined whether similar modulation is also present in non-cholinergic VP neurons. We selectively expressed GCaMP in ChAT-negative neurons following delivery of a Cre-Off AAV [41,45] into the VP of ChAT-IRES-Cre mice and performed analogous FP recordings **(Fig 5A and Fig S3)**. We first monitored non-cholinergic VP neuron activity over four days of CRF-1s training and found consistent positive responses to lever extension, lever press, and reward across days **(Fig. 5B,C and Fig. S4)**. In addition, we observed retrieval-dependent activation in response to reward **(Fig. 5B)** but not lever extension or press **(Fig. S4C, D)**. Next, we assessed whether non-cholinergic VP neuron activity was modulated by reward dilution.

**Figure 5.**
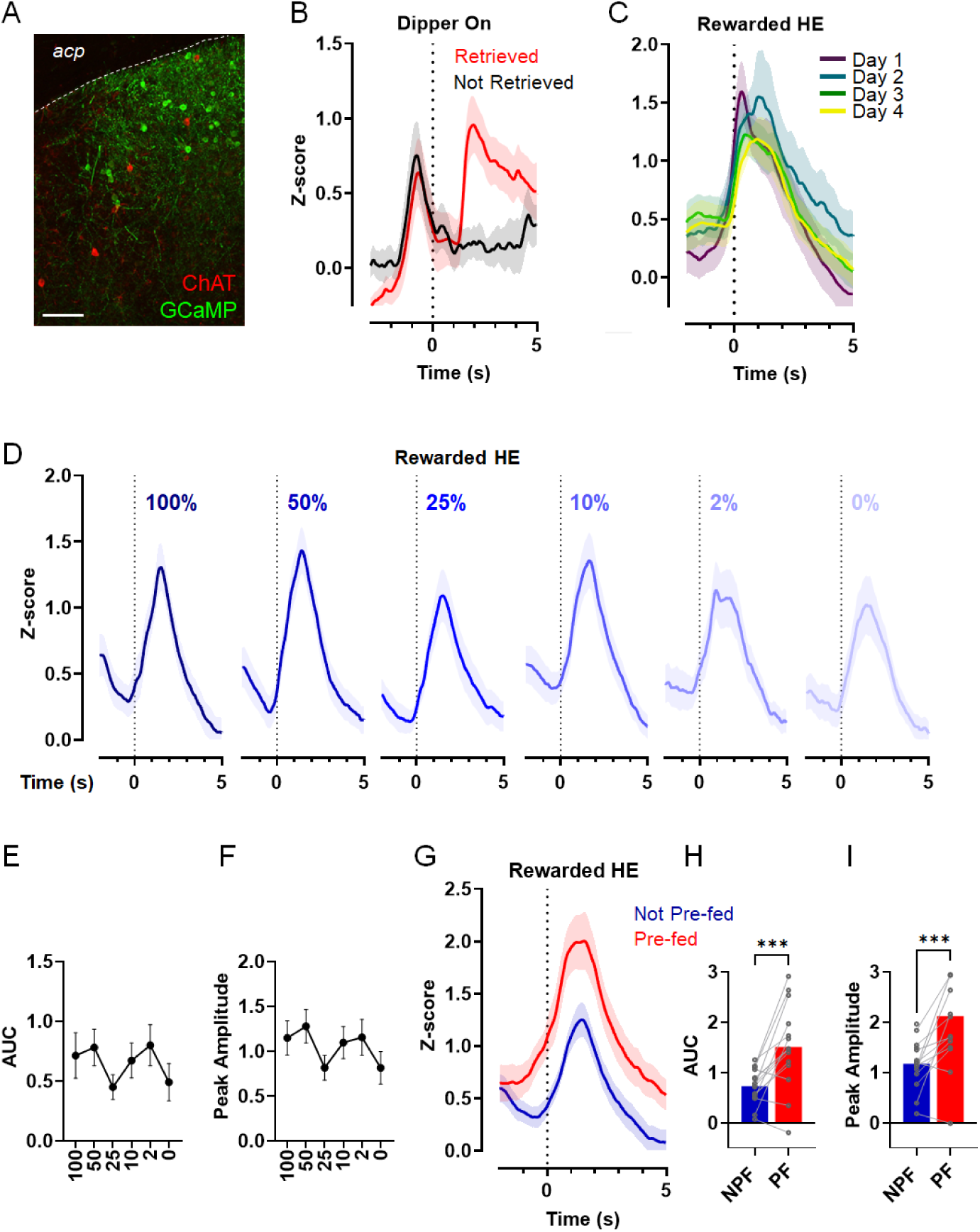
Non-cholinergic VP neuron reward responses are sensitive to satiety but not reward palatability. **A.** Co-immunolabeling of the Cre-Off FAS-GCaMP6f and ChAT revealed a lack of co-localization in VP of a ChAT-IRES-Cre mouse 4 weeks after injection. Scale bar, 100 μm. *acp* = anterior commissure. **B.** Mean z-scored GCaMP fluorescence aligned to Dipper On measured on the 4^th^ training day, showing a distinct increase in activity in retrieved vs not-retrieved trials. Note that the preceding lever press activity is independent of retrieval. **C.** Mean GCaMP activity across training days aligned to the first Rewarded HE in each trial, showing no effect of training. **D.** Mean z-scored GCAMP signals time-locked to Rewarded HE at each milk concentration. **E, F.** Neither A.U.C. (F _(2.237, 30.88)_ = 1.574, p = 0.2219) nor peak amplitude (F _(1.962, 27.07)_ = 1.185, p = 0.3203) were significantly altered by milk dilution. **G.** Mean z-scored GCAMP responses time-locked to Rewarded HE in NPF vs PF sessions. **H, I.** Rewarded HE response A.U.C. and peak amplitude were significantly increased following PF (***p < 0.001). Data shown as mean ± SEM. n = 15 mice.

Despite causing a clear reduction in behavioral performance in these mice **(Fig. S5)**, reward dilution did not significantly alter neural responses to reward retrieval **(Fig. 5D-F)** or lever press **(Fig S6D, E)**. However, we found a significant effect of reward dilution on the response to lever extension **(Fig. S6A-C)**, suggesting that in non-cholinergic VP neurons, cue-evoked responses are sensitive to reward palatability.

Lastly, we tested whether PF-induced satiety affects non-cholinergic VP neuron reward-evoked activity. In these neurons, PF led to a robust increase in the reward-evoked response compared to the NPF control condition **(Fig 5G-I)**, but no effect on peak lever extension or press responses **(Fig. S7)**. These responses are reminiscent of those seen in VP-CNs, suggesting that reward-evoked activity in the VP is broadly influenced by satiety.

## DISCUSSION

Using cell-targeted fiber photometry recordings, we have revealed that VP-CN activity increases in response to cues, actions, and reward retrieval in a reinforcement task. These responses were consistent across sessions and not altered by behavioral training. The unconditioned reward evoked the most robust activation, a response that was uniquely sensitive to two different reward devaluation conditions. We found that reward-evoked VP-CN activity was decreased as rewards were diluted. In contrast, VP-CN responses to the reward were enhanced when mice were pre-fed the reward prior to the task. These results suggest that VP-CN activity is not only recruited by reward retrieval/consumption, but that it is also dynamically modulated by reward palatability and physiological state.

Single-unit recording studies have reported that most VP neurons respond to palatable reward delivery and consumption, and that they do so primarily via phasic excitation [13,16,17]. The observed responses to the unconditioned stimulus (UCS) were present early on and remained unchanged with training [13], suggesting that they are related to stable sensory properties of the UCS. Yet these responses have also been shown to be reward-selective and can track relative hedonic value or size [14–17]. While the identity of the neuron types in those studies was not verified, more recent work has suggested a prominent role for GABAergic neurons [32,33]. Our data, which shows similar training-independent and value-sensitive reward-evoked activity in VP-CNs, suggests that the VP’s dynamic encoding of reward information involves cholinergic transmission.

Ample evidence links VP activation to feeding behavior and preferred reward choices [46–48]. While the specific cellular sources driving VP-CN activation in response to reward remain to be determined, these may include excitatory inputs from PFC, BLA, and lateral hypothalamus, which have been shown to target the VP and are known for roles in reward learning, decision-making and reward-seeking [3,5]. Alternatively, this reward-evoked activation could be indirectly mediated by a reduction in GABAergic inhibition by NAc inputs [16]. Indeed, single-unit recordings have shown that most reward-responsive NAc neurons in rats are inhibited by sucrose reward consumption [49–51]. This is in line with more recent work demonstrating that both medium spiny neuron (MSN) subtypes show decreased Ca^2+^ activity during ongoing consumption of sweet milk in mice [52], which could enable a phasic disinhibition of VP-CNs.

Interestingly, we found that the magnitude of VP-CN reward-evoked activation is diminished by reward dilution. This finding suggests that, beyond signaling the retrieval of the reward, VP-CN population activity provides dynamic information about palatability. In contrast, in non-cholinergic neurons, dilution did not significantly reduce phasic responses to the reward, revealing cell type-specific differences in how palatability influences reward-evoked activity in the VP. However, we cannot rule that the non-cholinergic population-level signal reflects heterogeneous contributions from distinct GABAergic and glutamatergic subpopulations.

By establishing synaptic contact with VP-CNs [53], NAc afferents provide a direct neuroanatomical route by which changes in reward palatability could influence VP-CN activity. Consistent with a role for the NAc in hedonic processing, enhancement of sucrose “liking” reactions requires simultaneous opioid activation in portions of the NAc medial shell and the posterior VP [54]. Further, electrophysiological recordings in the rat NAc shell and core have identified a small fraction of NAc neurons that are sensitive to changes in sucrose concentration. Of these, 75% displayed increased firing rates at higher sucrose concentrations [51]. Therefore, at the diluted milk concentrations used in our study we would have predicted reduced reward activity in these palatability-coding NAc neurons and a concomitant increase in VP-CN activity — the opposite of what we observed. However, it is still possible that reduced NAc activity in response to diluted rewards could disinhibit a yet to be determined intermediate signal that inhibits VP-CNs. Nonetheless, recent evidence has also identified peripheral and central gustatory innervation of NAc shell D1-MSNs [55], raising the intriguing possibility of a gustatory-NAc-VP circuit shaping VP-CN palatability-related responses.

It is important to note VP neuronal activity in response to preferred palatable rewards may precede that of the NAc upon reward delivery [17]. Therefore, we speculate that VP-CN responses to changes in palatability are unlikely to be dictated solely by upstream NAc GABAergic input. For example, reward-responses in VP-projecting VTA GABA neurons have been shown to correlate with reward palatability [56], which could in turn influence palatability encoding in VP-CNs. Other candidate mechanisms implicated in encoding reward value include attenuation of glutamatergic inputs to the VP from regions such as the BLA, or reduced modulation by endogenous opioids and orexin [57–60]. An important objective for future work will be to identify the neuroanatomical and cellular mechanisms behind the sensitivity of VP-CNs to this form of reward devaluation.

Unlike the diminished VP-CN reward response seen following reward dilution, our results reveal that pre-feeding mice with the reward *increased* VP-CN reward-evoked activity in the task. These findings suggest that VP-CNs activate more strongly to rewards during satiety compared to food-restriction. Further, this pattern contrasts with that observed following reward dilution, indicating that VP-CN activity is differentially sensitive to distinct forms of reward devaluation. Because both manipulations reduced behavioral responding, the increased reward-evoked GCaMP response after pre-feeding is unlikely to be explained by differences in pressing or retrieval behavior relative to the non-prefed condition. While we cannot rule out potential differences in licking microstructure following pre-feeding versus dilution conditions, both satiety and reduced palatability have been associated with reduced licking behavior [30,61,62]. We also show that pre-feeding did not significantly alter phasic responses to lever extensions and lever presses, suggesting that satiety does not lead to a generalized increase in task-evoked VP-CN activity but that it is relatively specific to reward retrieval.

The increased VP-CN phasic reward response following pre-feeding may reflect greater or more sustained firing during reward retrieval of VP-CNs already activated by reward. Alternatively, the enhanced response could originate from a different subset of VP-CNs that only becomes engaged following satiety, boosting overall population activity. Indeed, the notion that VP-CNs can be functionally heterogeneous is supported by recent identification of two VP-CN subpopulations that distinctly respond to either appetitive or aversive olfactory stimuli [36].

Notably, non-cholinergic VP neurons also showed enhanced reward-evoked activity after pre-feeding, pointing to a shared upstream mechanism that broadly activates different VP neuron populations. While local GABA collateral activity may regulate intra-VP activity [63], this is unlikely to be a main driver of the *increased* neuronal activity seen with pre-feeding. Instead, a decrease in GABAergic input from NAc MSNs to the VP represents a compelling candidate mechanism, aligning with evidence that satiation suppresses activity in both NAc MSNs and VTA dopamine neurons [64–67]. Other satiety-sensitive inputs, such as those from the lateral hypothalamus [3,68], could also influence VP activity directly or indirectly and warrant further investigation.

Our findings demonstrate that VP-CNs are primarily activated by reward, and that these responses are dynamically shaped by reinforcer properties and satiety. These results expand our current understanding of VP function in reward-related behavior by identifying VP-CNs as distinct players in value- and state-dependent reward processing. Future strategies that specifically target VP cholinergic function may provide further insight into reward processing and its dysfunction in neuropsychiatric disorders.

## ACKNOWLEDGMENTS

We thank Christoph Kellendonk for sharing the Cre-Off-GCaMP viral vector and Talia Lerner and Venus Sherathiya for their support with Guppy software for fiber photometry data analysis.

## AUTHOR CONTRIBUTIONS

A.G., R.S., C.M.M., R.E.S., D.R.Z., J.E.B. and E.F.G. conducted experiments and data analysis. E.F.G. and A.G. wrote the manuscript. E.F.G. designed the experiments. E.F.G. supervised the experiments and data analysis.

## FUNDING

This work was supported by NIDA R01-DA055018 to E.F.G. and Len Blavatnik STEM Research Fellowships and Fordham Undergraduate Research Grants to R.S. and A.G.

## COMPETING INTERESTS

The authors have nothing to disclose.

## Notes

### Competing Interest Statement

The authors have declared no competing interest.

