## Supplementary Figures for "Ventral Pallidum Cholinergic Neurons Respond to Reward and Signal Reward Value"

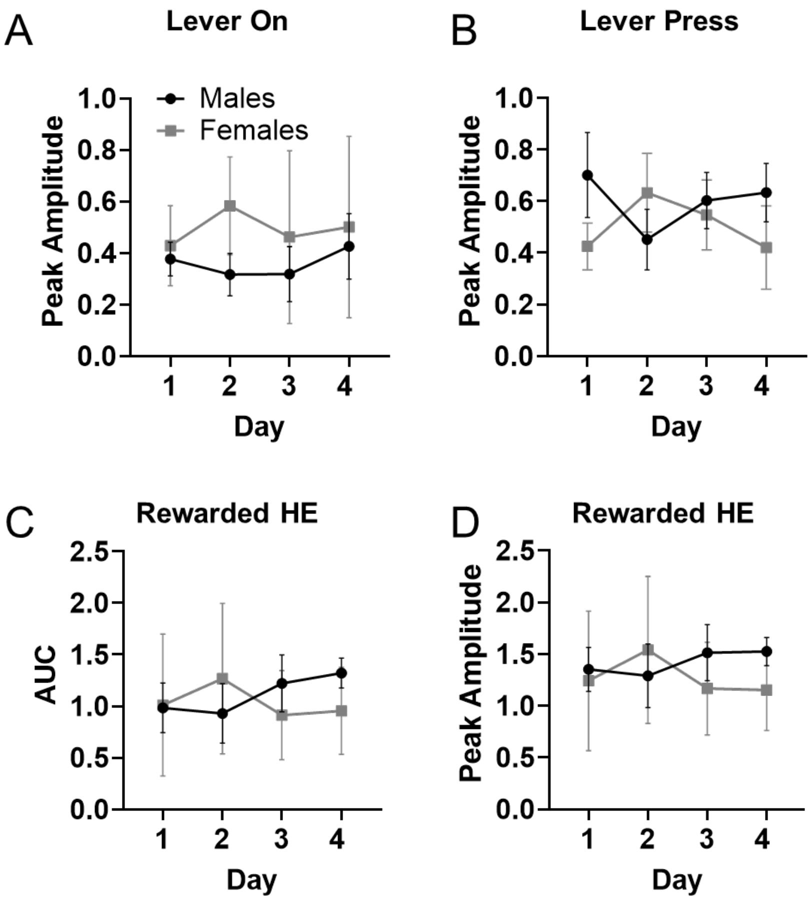


**Figure S1. Analysis of VP-CN responding across CRF training days by sex.** Mean GCaMP activity in VP-CNs of males and females aligned to Lever ON (A), Lever Press (B) and the Rewarded Head Entry (C, D). No significant main effects of sex or training day were found across all measures using 2-way RM ANOVA, $p > 0.05$. Data shown as mean ± SEM. n = 12 males, 6 females.

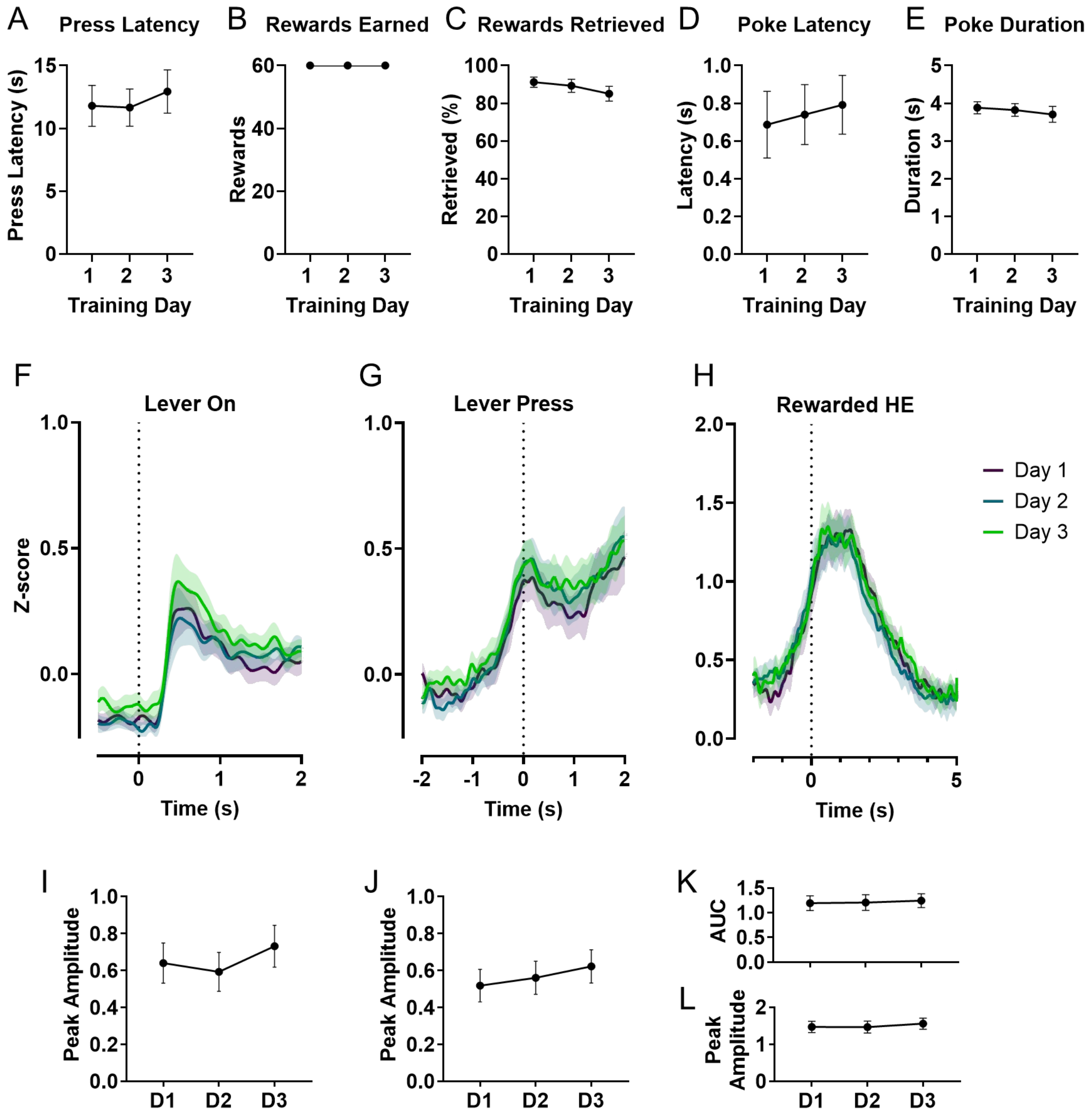


**Figure S2. VP-CN reward-related activity is stable across CRF-2s sessions. A-E.** Behavioral performance in the CRF-2s task, where reward was delivered 2 seconds after the press, did not change significantly across 3 days. **F-H**. Mean z-scored GCaMP fluorescence aligned to Lever On, lever press, and Rewarded HE across the same 3 days. Dotted line denotes behavior event onset. Peak amplitude was not altered by training day when aligned to Lever ON (**I**), $F_{(1.807, 30.72)} = 2.307$, $p = 0.1210$ or Lever Press (**J**), $F_{(1.872, 31.82)} = 1.451$, $p = 0.2491$). **K, L.** Neither VP-CN activity A.U.C. ($p = 0.7979$) or peak amplitude ($p = 0.1638$) in response to Rewarded HE were affected by training. Data shown as mean ± SEM. n = 18 mice.

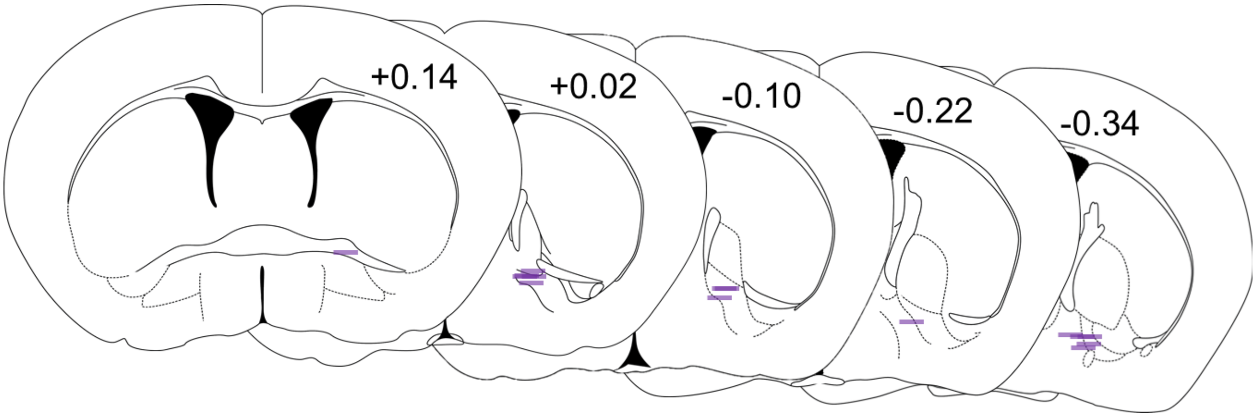


**Figure S3. Unilateral fiber optic cannula placements for GCaMP recordings from non-cholinergic VP neurons.** Summary of fiber optic cannula placements for experimental mice (n = 15), projected onto schematic coronal sections from the Paxinos and Franklin mouse brain atlas (3rd ed). Purple lines represent the estimated location of individual fiber tips, identified as the ventral-most point of the fiber track across serial histological sections. Numbers indicate distance (mm) from Bregma.

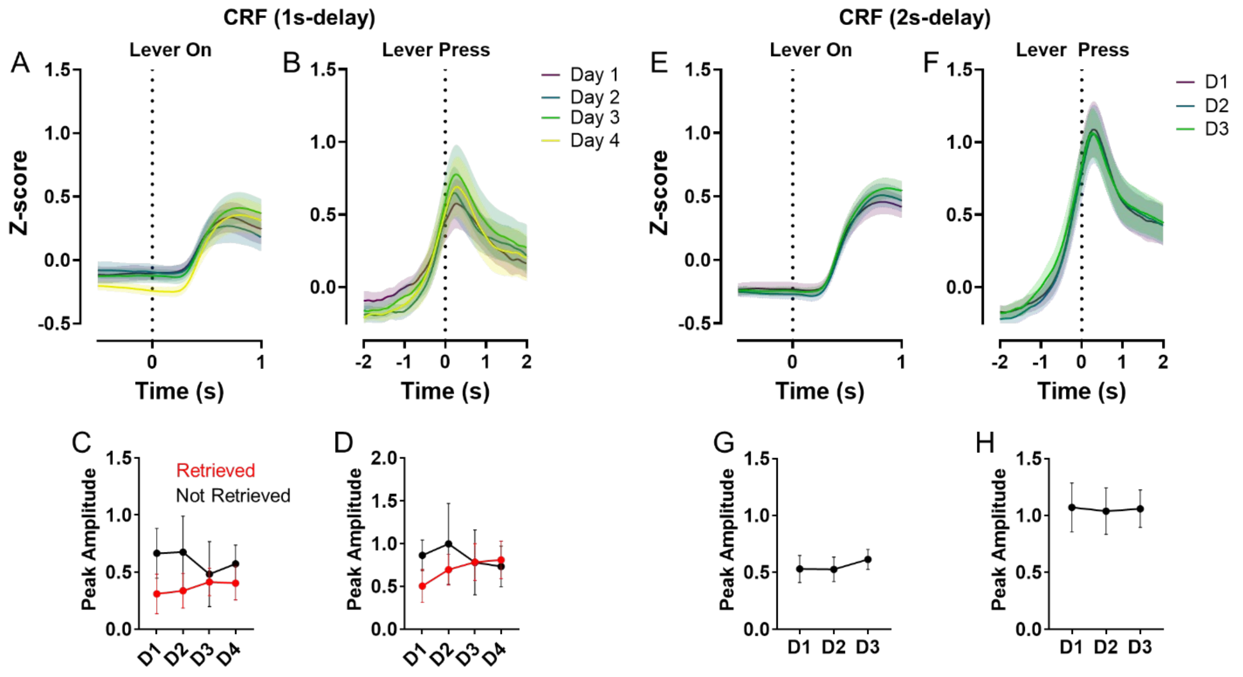


**Figure S4. GCaMP responses of non-cholinergic VP neurons to lever extension and lever press are stable across CRF training days.** Cre-off GCaMP recordings from ChAT-IRES-Cre mice revealed increased phasic activity in response to LeverOn (**A**) and Lever Press (**B**) in the CRF-1s task. **C**. The increase following Lever On was similar across CRF-1s days and was independent of reward retrieval. **D.** Lever Press responses were not affected by training or retrieval. Similarly, Lever On (**E, G**) and Lever Press (**F, H)** responses were stable across CRF-2s days. Data shown as mean ± SEM. n = 15 mice.

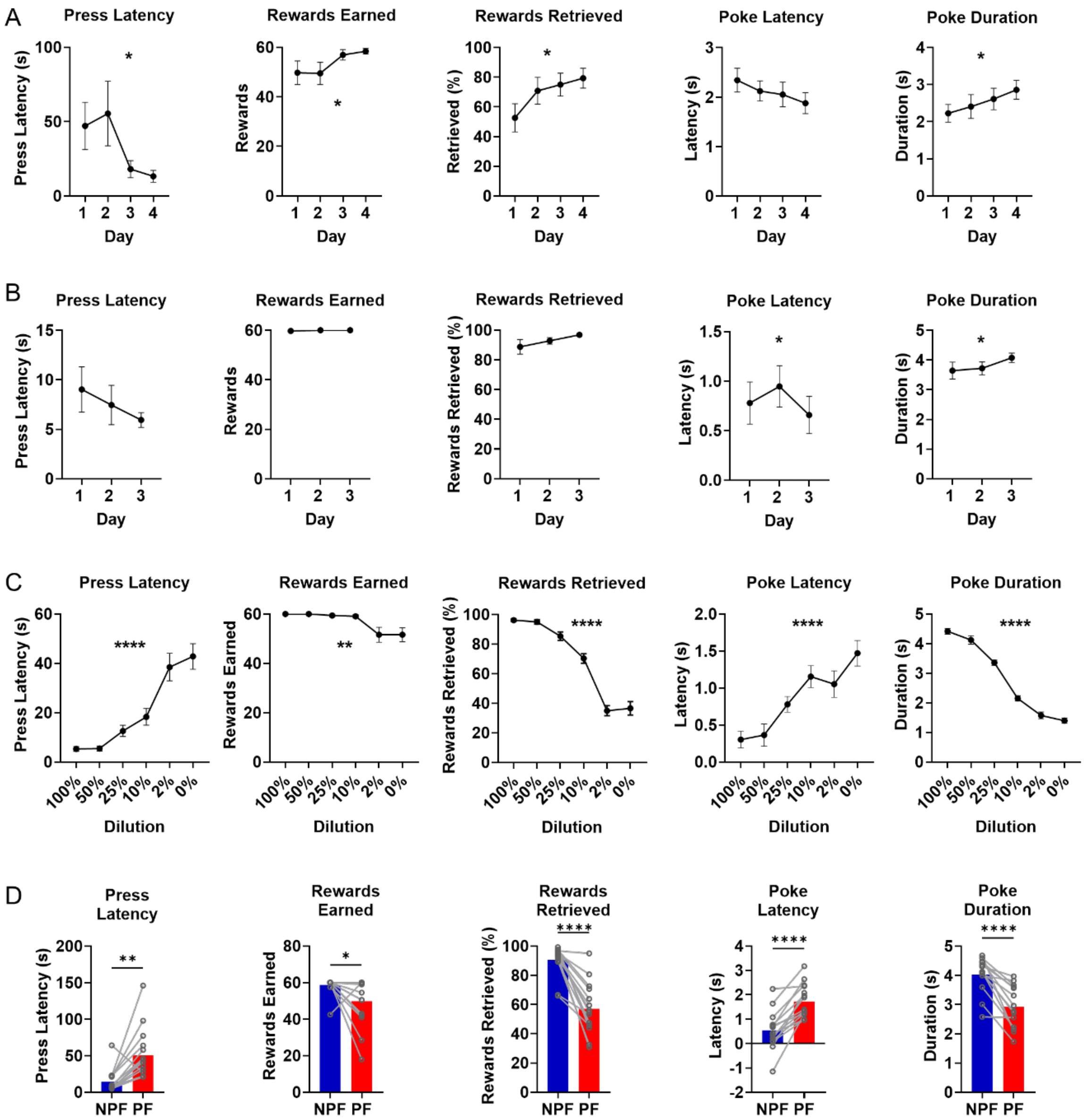


**Figure S5. Behavioral analysis of CRF performance for mice expressing GCaMP in non-cholinergic VP neurons. A.** In CRF-1s, mice showed improved task performance over days, including increased rewards earned and retrieved, and greater time spent in the feeder port during reward presentation. Press latency was also significantly reduced *p <0.05). **B**. In the CRF-2s task, poke latency ($F_{(1.918, 26.85)} = 4.196$, $p = 0.0273$) and duration $F_{(1.512, 21.16)} = 6.893$, $p = 0.0084$) showed a significant effect of day, which was not observed on press latency, rewards earned or retrieved. **C.** Behavioral performance across CRF-2s sessions as a function of milk reward concentration. RM one-way ANOVA indicated a significant effect of dilution on all measures. ** p<0.01, ****p <0.0001. **D**. Two-day average performance in CRF-2s sessions with and without pre-feeding with the milk reward (PF vs NPF). Paired t-tests revealed significant increases in press latency and poke latency, and significant reductions in rewards earned, rewards retrieved, and in poke duration (*p < 0.05, **p < 0.01; ****p<0.0001). Data shown as mean ± SEM. n = 15 mice.

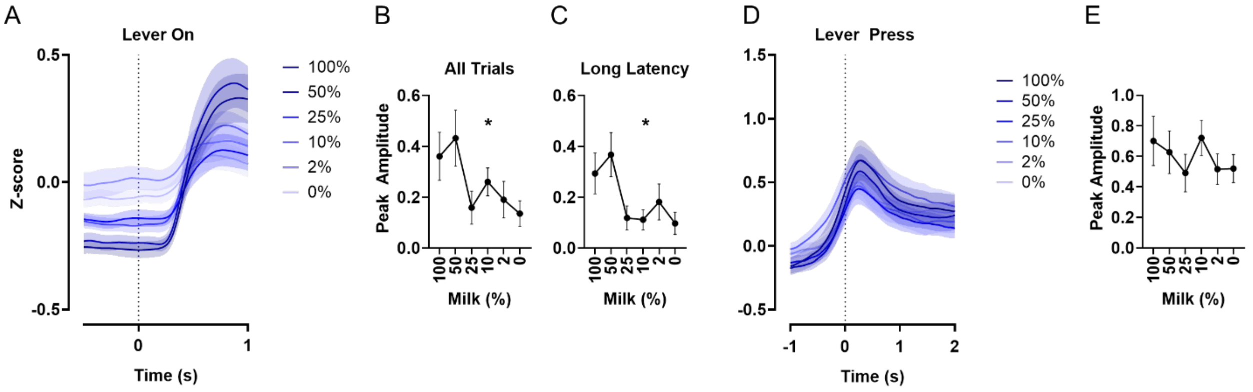


**Figure S6. Effect of reward dilution on GCaMP responses of non-cholinergic VP neurons to lever extension and lever press. A, B.** Mean peak amplitude of GCaMP responses aligned to Lever ON was significantly reduced by dilution ($F_{(2.166, 30.33)} = 4.844$, $p = 0.0131$). **C.** A similar effect was observed in Lever On trials where the press latency was longer than 3 s ($F_{(2.477, 34.68)} = 3.408$, $p = 0.0356$) **D, E.** Lever press-related activity was not significantly altered by reward dilution ($F_{(1.687, 23.62)} = 1.991$, $p = 0.1638$). Data shown as mean ± SEM. n = 15 mice.

.

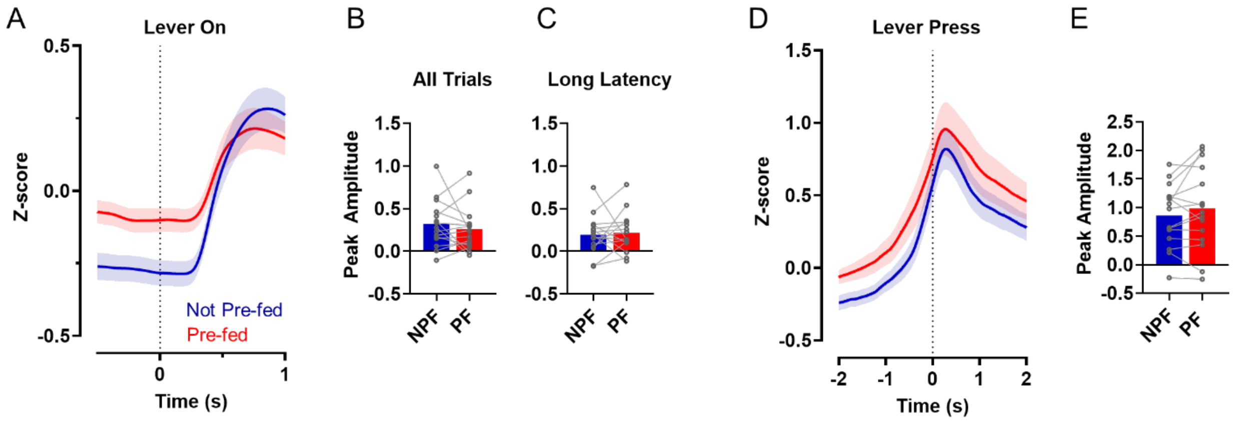


**Figure S7. Effect of pre-feeding on GCaMP responses of non-cholinergic VP neurons to lever extension and lever press.** No significant differences were observed following PF in GCaMP responses to Lever On (**A-C**) or to Lever Press (**D, E**), $p > 0.05$. Data shown as mean ± SEM. n = 15 mice.
